# Determining the content of vesicles captured by golgin tethers using LOPIT-DC

**DOI:** 10.1101/841965

**Authors:** John J.H. Shin, Oliver M. Crook, Alicia Borgeaud, Jérôme Cattin-Ortolá, Sew-Yeu Peak-Chew, Jessica Chadwick, Kathryn S. Lilley, Sean Munro

**Affiliations:** MRC Laboratory of Molecular Biology, Francis Crick Avenue, Cambridge, CB2 0QH, UK; The Milner Therapeutics Institute, University of Cambridge, Puddicombe Way, Cambridge, CB2 0AW, UK; Cambridge Centre for Proteomics, Department of Biochemistry, University of Cambridge, Cambridge, UK; MRC Biostatistics Unit, Cambridge Institute for Public Health, Cambridge, UK

## Abstract

The internal organisation of the cell depends on tethers at destination organelles to selectively capture incoming transport vesicles to facilitate SNARE-mediated fusion. The golgin long coiled-coil proteins function as tethers that contributes to this specificity at the Golgi (1). Golgin-97, golgin-245 and GCC88 golgins of the *trans*-Golgi capture vesicles derived from endosomes, which serve to recycle the critical Golgi machinery required to deliver lysosomal hydrolases and to maintain exocytosis. Retrograde trafficking from endosomes to the *trans*-Golgi network (TGN) is a complex process that involves the sorting of transmembrane cargo proteins into distinct transport vesicles by adaptors from multiple pathways. The content of these distinct vesicles, which golgin they target and the factors that mediate this targeting are not well understood. The major challenges that have limited advances in these areas is the transient nature of vesicle tethering, and the redundancies in their mechanisms that confound experimental dissection. To gain better insight into these problems, we performed organelle proteomics using the Localisation of Organelle Proteins by Isotope Tagging after Differential ultraCentrifugation (LOPIT-DC) method on a system in which an ectopic golgin causes vesicles to accumulate in a tethered state (2). By incorporating Bayesian statistical modelling into our analysis (3), we determined that over 45 transmembrane proteins and 51 peripheral membrane proteins of the endosomal network are on vesicles captured by golgin-97, including known cargo and components of the clathrin/AP-1, retromer-dependent and -independent transport pathways. We also determined a distinct class of vesicles shared by golgin-97, golgin-245 and GCC88 that is enriched in TMEM87A, a multi-pass transmembrane protein of unknown function that has previously been implicated in endosome-to-Golgi retrograde transport (4). Finally, we categorically demonstrate that the vesicles that these golgins capture are retrograde transport vesicles based on the lack of enrichment of lysosomal hydrolases in our LOPIT-DC data, and from correlative light electron tomography images of spherical vesicles captured by golgin-97. Together, our study demonstrates the power of combining LOPIT-DC with Bayesian statistical analysis in interrogating the dynamic spatial movement of proteins in transport vesicles.

## Introduction

A fundamental question in membrane traffic is how transport vesicles recognise the correct compartment with which to fuse (5). The Golgi apparatus is an ideal system to study this issue as it functions as a central sorting hub for membrane traffic that sends and receives transport vesicles to and from numerous destinations. The golgins, a large and ancient family of ubiquitously expressed long coiled-coil proteins of the Golgi, function as tethers that contribute to the specificity of membrane traffic by selectively capturing incoming transport vesicles from destination organelles (1). However, the molecular players that allow a specific vesicle to connect to a specific golgin, and the content that each specific vesicle delivers remains poorly defined.

Each Golgi compartment is decorated with a distinct set of golgins that are anchored to the membrane via their C termini, with golgin-97, golgin-245, GCC88 and GCC185 at the *trans*-Golgi in mammalian cells. Mutation of these *trans*-Golgi golgins result in only mild phenotypes, suggesting redundancies between them that have made it challenging to distinguish their functions. An in vivo assay for golgin tethering activity overcame this problem by testing whether the golgins are sufficient, rather than necessary, to function as Golgi tethers (1). Golgins were relocated to an ectopic site by swapping their C termini with a mitochondrial transmembrane domain (golgin-mito), which was sufficient to redirect specific transport vesicles to the mitochondria and keep them in a permanently tethered state (Fig. 1A). Golgin-97-mito, golgin-245-mito and GCC88-mito specifically captures endosome-to-Golgi vesicles. Whereas, GCC185 was not able to confer vesicle capture activity using this assay, but has been implicated as a Golgi tether using alternative systems (6). Golgin-97 and golgin-245 share a conserved vesicle binding domain which they use to capture the same pool of vesicles (7). These domains bind directly to TBC1D23, a member of a family of Rab GTPase-activating proteins, which bridges golgin-97 and golgin-245 to endosome-to-Golgi vesicles and the WDR11 complex (Fig. 1A) (8).

**Figure 1.**
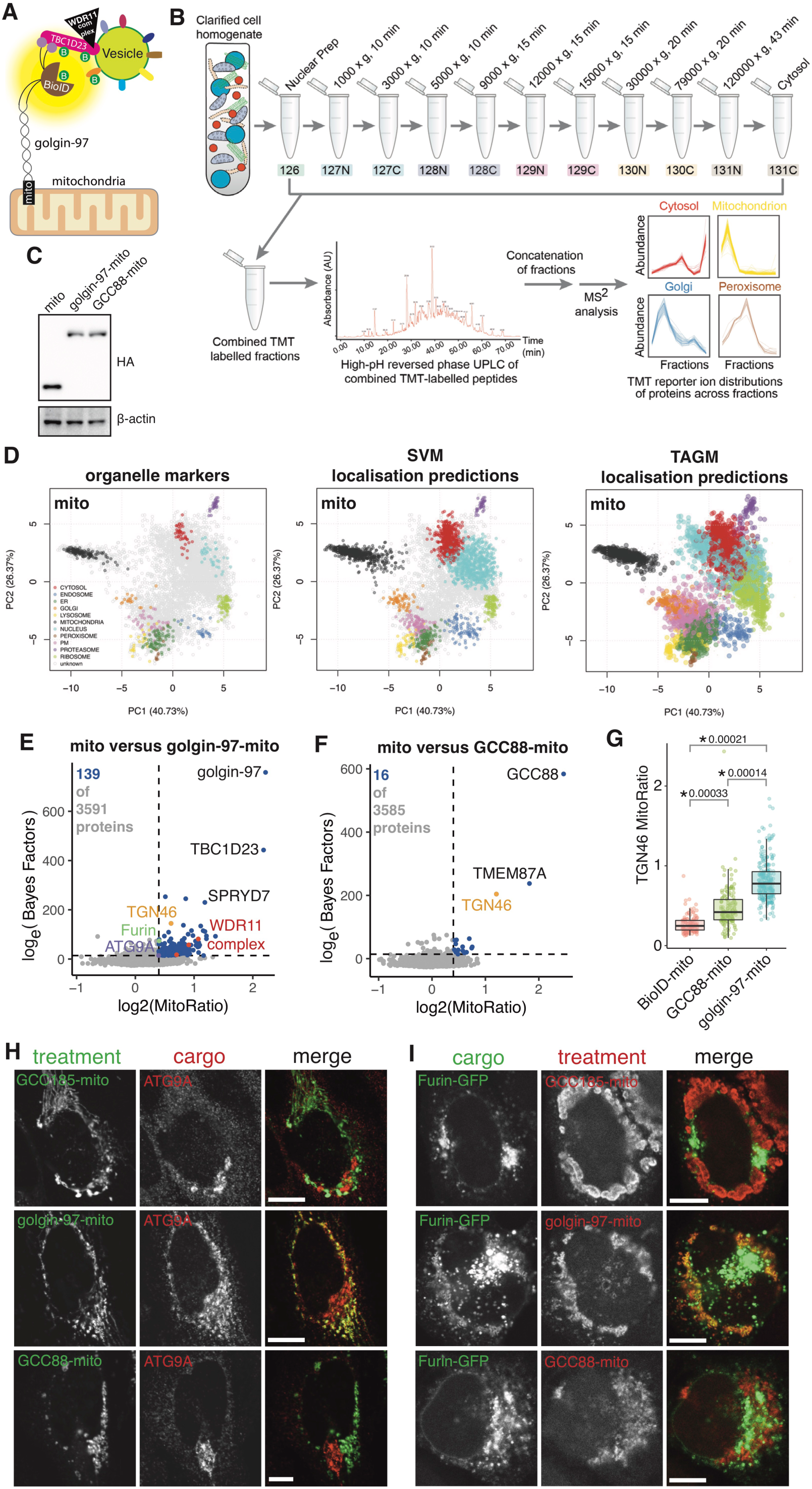
LOPIT-DC on mitochondrial relocated golgin-97 and GCC88. **A**. Applying proximity biotinylation to mitochondrial relocated golgin-97 and golgin-245 identifies their interaction with TBC1D23 and the WDR11 complex on endosome-to-Golgi vesicles. **B**. Overview of the LOPIT-DC workflow. Cells are gently lysed and fractionated through differential ultracentrifugation. The peptides of each fraction are then multiplexed with 11-plex TMT, pre-fractionated through high-pH reversed phase UPLC and then analysed by MS^2^ or SPS-MS^3^. This gives the profiles of the proteome of the cell, which is then analysed in (D). **C**. Immunoblot of proteins from 293T cells stably expressing the doxycycline-inducible mito, golgin-97-mito and GCC88-mito with 1 *µ*g ml^−1^ doxycycline for 48 h. **D**. Principle component analysis (PCA) projections for the LOPIT-DC showing organelle markers before classification, after SVM classification and after TAGM classification. **E,F**. Results of MitoRatio versus Bayes Factor analysis comparing mito to golgin-97-mito and GCC88-mito. Thresholds are based on experimentally confirmed hits (see text) (log_2_(MitoRatio) ≥ 0.4, log_e_(Bayes Factor) ≥ 14). Mito versus golgin-97-mito: 139 proteins not including golgin-97 were identified within threshold of 5914 analysed proteins (3591 proteins after pre-filtering). Mito versus GCC88-mito: 16 proteins not including GCC88 were identified within threshold of 5532 analysed proteins (3585 proteins after pre-filtering). **G**. Quantification of confocal micrographs of 293T cells used in LOPIT-DC (Fig. S2A) measuring the ratio of the mean intensity of TGN46 at the mitochondria over its intensity at the Golgi (MitoRatio). Boxplots are of MitoRatios of all cells quantified across 3 independent replicates. p-values are of two-sample t-tests comparing the overall mean MitoRatio of each replicate. A minimum of 115 cells were quantified per treatment. **H,I**. Confocal micrographs of HeLa cells expressing the indicated golgin-mito construct (HA) stained for endogenous ATG9A or co-transfected with Furin-GFP. Scale bars, 10 *µ*m.

Many open questions regarding the *trans*-Golgi golgins remain. Golgin-97 and golgin-245 have been reported to bind AP-1 derived vesicles through TBC1D23 and possibly the WDR11 complex; however, it is evident that AP-1 independent vesicles are also captured by these golgins (9). Moreover, TBC1D23 is expressed only in metazoans, yet golgin-97 is conserved as IMH1 in yeast (10). Therefore, an ancestral population of vesicles that connect to golgin-97 and golgin-245 using a TBC1D23-independent mechanism is likely to exist which remains to be characterised. Finally, the vesicle-associated specificity factors that target transport vesicles to GCC88 are unknown, and the physiological roles that distinguish GCC88 from golgin-97 and golgin-245 are still poorly defined. To elucidate these questions, we set out to determine the content of the vesicles that are specifically captured by golgin-97 and GCC88. Given the subtle phenotypes of golgin mutants and the possibility of these mutants adjusting over time by switching on compensatory pathways, we refrained from taking a genetic approach. Instead, we opted to combine the mitochondrial relocation assay with proteomics to determine which vesicle cargo, adaptors and accessory proteins are redirected to the mitochondria by golgin-97-mito and GCC88-mito.

Applying proximity biotinylation to mitochondrial relocated golgins have been successfully used in the past to identify novel protein interactions (Fig. 1A) (8). However, its limited biotinylation range prevents this approach from determining the entire content of vesicles. Moreover, it is also skewed against the identification of small proteins and proteins that may have a limited amount of exposed lysines to biotinylate, such as in the short tails of tail-anchored transmembrane proteins which are often cargo found on vesicles. In addition, mitochondrial purification by cell fractionation and immune-purification each come with their own caveats. The former does not ensure the complete separation of organelles and does not account for the change in size and density of the mitochondria due to vesicle capture across treatments. The latter depends heavily on the specificity of the antibody, and its stringency may lead to the loss of important transient interactions involved in tethering.

LOPIT-DC combines cell fractionation with quantitative mass spectrometry and multivariate data analysis to allow the high-throughput and simultaneous characterisation of multiple subcellular compartments, without the requirement for total purification of compartments of interest (Fig. 1B,D) (2). To avoid the issues of alternative approaches, we utilised LOPIT-DC to determine the content of vesicles captured by golgin-97-mito and GCC88-mito (2).

## Results and Discussion

### LOPIT-DC on mitochondrial relocated golgin-97 and GCC88

An overview of the LOPIT-DC workflow we used is shown in Fig. 1B,D. LOPIT-DC was applied to HEK 293T cells expressing golgin-97-mito, GCC88-mito or a mitochondrial transmembrane domain (mito) as a control (Fig. 1C). The profiles of approximately 6000 proteins analysed by conventional tandem mass spectrometry (MS^2^) present across three independent replicates for each treatment were obtained. A prediction of the subcellular localisation for each protein was then made by matching to profiles of known organelle markers using supervised machine learning through the support vector machines (SVM), and using Bayesian statistical modelling through the T-Augmented Gaussian Mixture model (TAGM) method (Fig. 1D) (3). The latter takes into account the uncertainty that arises when classifying proteins that reside in multiple locations, or unknown functional compartments and also those that dynamically move within the cell, providing a richer overall analysis of our spatial proteomics data (3).

The performance of tandem mass tag (TMT)-based quantification by MS^2^ can be affected by interference from contaminant peptides with similar properties to the target peptide, which is resolved with an additional round of ion selection and fragmentation in synchronous precursor selection mass spectrometry (SPS-MS^3^) (11). Our LOPIT-DC data analysed by MS^2^ showed effective separation of organelles with good resolution (Fig. S1A,B), even though the overall resolution of SPS-MS^3^ is somewhat better (2). However, our application of LOPIT-DC was not designed to resolve the subtle differences in the overall profiles of different organelles in the cell, but instead to observe dramatic changes in the profile of proteins across treatments, a requirement that was satisfied by our MS^2^ analysis.

To identify proteins that translocate to the mitochondria as a result of our mitochondrial golgins, we first leveraged the rich spatial information of our LOPIT-DC data by applying an independent pre-filtering step. Proteins predicted by TAGM to localise to the mitochondria or nucleus in the mito control were discarded from subsequent analysis. These proteins are likely resident mitochondrial and nuclear proteins that are not relevant to our biological question. Discarding them made it easier to discern proteins that shift toward a mitochondrial profile in golgin-mito cell lines (Fig. S1C).

We next analysed our data by quantifying general perturbations in protein profiles between golgin-mito cells and control cells using a Bayesian non-parametric two sample test (Bayes Factor), and by quantifying for protein profiles that shift towards a mitochondrial profile by using a mitochondrial ratio (MitoRatio) (see methods). As expected, our strongest hits when comparing our LOPIT-DC of golgin-97-mito to the mito control was golgin-97 itself and TBC1D23 (Fig. 1E). Likewise, GCC88 was the strongest hit when comparing GCC88-mito to mito (Fig. 1F). Strikingly, golgin-97-mito had more hits than GCC88-mito, suggesting that Golgin-97 had a greater overall ability to capture vesicles. This is consistent with our previous observations, and was further corroborated through quantitation of immunofluorescence microscopy (Fig. 1G & Fig. S2A) (7).

We next set a threshold for protein relocation to the mitochondria based on confirmed experimental observations (Fig. 1E,F). The major criterion was that this threshold should have the entirety of the WDR11 complex, composed of WDR11, FAM91A1 and C17orf75, as hits when comparing mito versus golgin-97-mito, but not when comparing mito versus GCC88-mito (Fig. 1A,E,F) (8). Application of this threshold also predicted ATG9A and Furin, normally resident in Golgi membranes and endosomes, as vesicle cargo specifically captured by golgin-97-mito (Fig. 1E) (12, 13). We verified this by immunofluorescence (Fig. 1H,I), demonstrating that the vesicles that recycle them to the Golgi can be captured by golgin-97 and golgin-245. Together, these data demonstrate the robustness of our Bayesian analysis of LOPIT-DC data.

### Cargo specific to golgin-97 and golgin-245

Proteins of the retrograde route to the TGN are in a constant state of flux as they are recycled throughout the endosomal network (14). Consistent with this phenomenon, we observed a strong enrichment for proteins of the endosomal network amongst those affected by golgin-97-mito. This includes proteins classified by our LOPIT-DC analysis and by multiple localisation databases as localised to endosomes, lysosomes, Golgi apparatus, plasma membrane (PM) or vesicles in our control. Of these, a total of 45 transmembrane proteins were robustly affected, including IGF2R (CIMPR), M6PR (CDMPR) and TGN46, which are known cargo of vesicles captured by golgin-97 and golgin-245 (Fig. 2A,B) (1, 7). In addition, we also detected 5 SNAREs involved in the fusion of endosome-to-Golgi vesicles at the *trans*-Golgi. Remarkably, the majority of these transmembrane proteins had profiles similar to endosomes in our control, and then shifted to a more mitochondrial profile as a result of golgin-97-mito (Fig. 2C,D). These shifts were partial, which is consistent with only the steady-state portion that localise to retrograde vesicles being relocated to the mitochondria. Based on these results, we define these transmembrane proteins as cargo of vesicles captured by golgin-97 and golgin-245.

**Figure 2.**
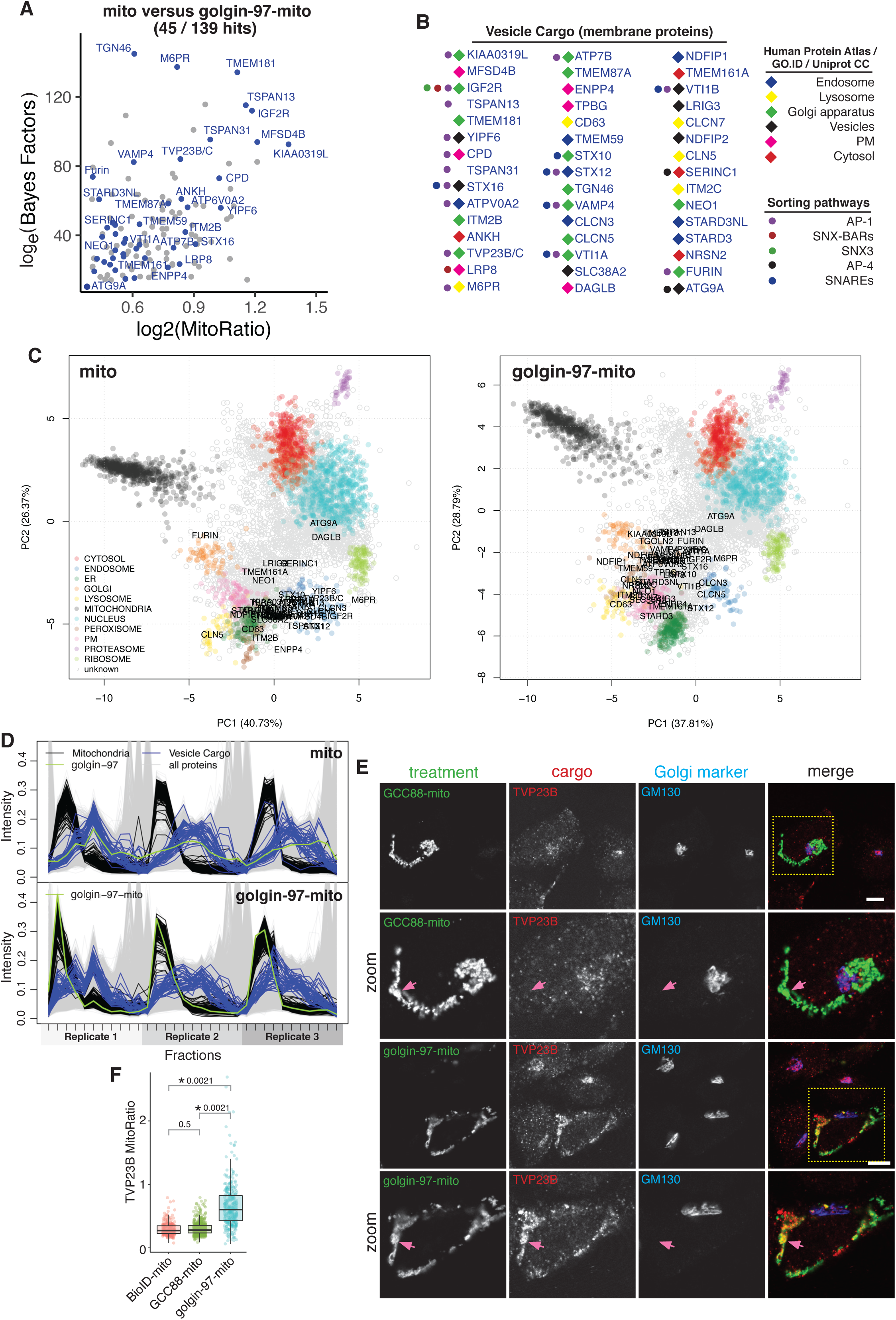
Cargo specific to golgin-97 and golgin-245. **A**. MitoRatio versus Bayes Factor analysis of mito versus golgin-97-mito showing all hits (blue) that are both localised to the endosomal network and are also transmembrane proteins. **B**. All hits shown in (A) ordered from left column to right column based on descending MitoRatio scores. Classification of their localisation and sorting pathway is based on multiple localisation databases and previous publications. **C**. PCA projections for the LOPIT-DC of mito versus golgin-97-mito showing SVM classifications and all hits from (A,B). **D**. TMT reporter ion distributions of proteins across fractions for each replicate for mito versus golgin-97-mito showing the profiles of known mitochondrial markers (Mitochondria) and all hits from (A,B) (Vesicle Cargo). **E**. Confocal micrographs of HeLa cells expressing the indicated golgin-mito construct (HA) stained for endogenous TVP23B (cargo) and GM130 (marker of *cis*-Golgi). **F**. Quantification of confocal micrographs of 293T cells used in LOPIT-DC (Fig. S2B) measuring the ratio of the mean intensity of TGN46 at the mitochondria over its intensity at the Golgi (MitoRatio). Boxplots are of MitoRatios of all cells quantified across 3 independent replicates. p-values are of two-sample t-tests comparing the overall mean MitoRatio of each replicate. A minimum of 209 cells were quantified per treatment. Scale bars, 10 *µ*m.

Retrograde trafficking to the *trans*-Golgi is typically challenging to interrogate because of the limited assays to determine the endosomal sorting pathways that are responsible for sorting specific cargo into specific vesicles. Of these different pathways, the content of clathrin/AP-1 derived vesicles is best resolved (15-17). We detected 16 AP-1 cargo that are affected by golgin-97-mito, which confirms previous reports of AP-1 vesicles binding to golgin-97 and golgin-245 (9). Of these, TVP23 (FAM18B) is a poorly characterised TGN protein that has been implicated in endosome to Golgi trafficking in yeast, secretion from the TGN in plants and plays a role in diabetic retinopathy in humans (18-20). We found that TVP23B is specifically relocated to the mitochondria by golgin-97-mito and not GCC88-mito (Fig. 2E,F & Fig. S2B). Furin is also a cargo of AP-1 that showed matching results (Fig. 1I). Together, this supports a conclusion that GCC88 has minimal ability to capture AP-1 derived vesicles.

A significant complicating factor in the study of retrograde trafficking is that many cargo proteins contain multiple sorting motifs that allow adaptors from numerous sorting pathways to bind to them. This redundancy has made it challenging to conclusively determine the necessity of some of these pathways. In addition to AP-1 cargo, we also identified 29 other cargo that are likely to be contained in transport vesicles derived from other sorting pathways which are captured by golgin-97 and golgin-245 (Fig. 2B). These proteins are thus candidates to be used as additional markers to further elucidate these pathways in the future. Such an example is with the autophagy protein ATG9A and its partner SERINC1, which are known to be sorted by AP-4 adaptors into vesicles secreted from the TGN to the periphery of the cell (21). Importantly, our analysis did not detect AP-4 or its accessory proteins, RUSC1 and RUSC2. This strongly suggests that ATG9A and SERINC1 are on retrograde vesicles derived from an, as yet, undefined sorting pathway. ATG9A is an important regulator of autophagosome formation and SERINC proteins have recently been identified as HIV restriction factors (13, 22, 23). Thus, their relocation to mitochondria by golgin-97-mito could provide a powerful assay to study the pathology of these proteins in the future.

### Adaptor proteins and accessory factors of vesicles specific to golgin-97 and golgin-245

The discovery of TBC1D23 and the WDR11 complex in Golgi tethering were derived from proximity biotinylation of vesicles captured by golgin-97-mito (8). This suggests that peripheral membrane proteins relocated to the mitochondria by golgin-97-mito are likely to be important adaptor proteins and accessory factors involved in the tethering process. A total of 51 peripheral membrane proteins of the endosomal network were affected by golgin-97-mito (Fig. 3A,B), including VPS45, an SM (Sec1p / Munc18) protein involved in the fusion of endosome-to-Golgi vesicles with the TGN. Strikingly, we found an enrichment for adaptor proteins and accessory factors from multiple endosomal sorting pathways. This included AP-1 adaptors (AP1G1, AP1S2, AP1AR, AP1M1 and AP1S1) and accessory proteins (CLINT1, SYNRG, PI4K2B and Rab9), but not their associated clathrin coat (15, 17, 24, 25). We also found proteins involved in retromer-dependent transport (SNX3, SNX4, VPS35, VPS26, VPS29), along with SNX-BAR proteins potentially involved in retromer-independent transport (SNX1, SNX2, SNX5 and SNX6) (26-28). Many peripheral membrane proteins are localised to multiple compartments in the cell, making them difficult to classify by spatial proteomics (Fig. 3C). However, we clearly observed the movement of these proteins to a more mitochondrial profile with golgin-97-mito (Fig. 3C-F). As with TBC1D23, we would expect that the more mitochondrial these peripheral membrane proteins become, the greater likelihood of them binding to golgin-97 (8). SPRYD7 is a protein of unknown function which localises endogenously to vesicles and fits this criteria (Fig. 1E & Fig. 3A) (29).

**Figure 3.**
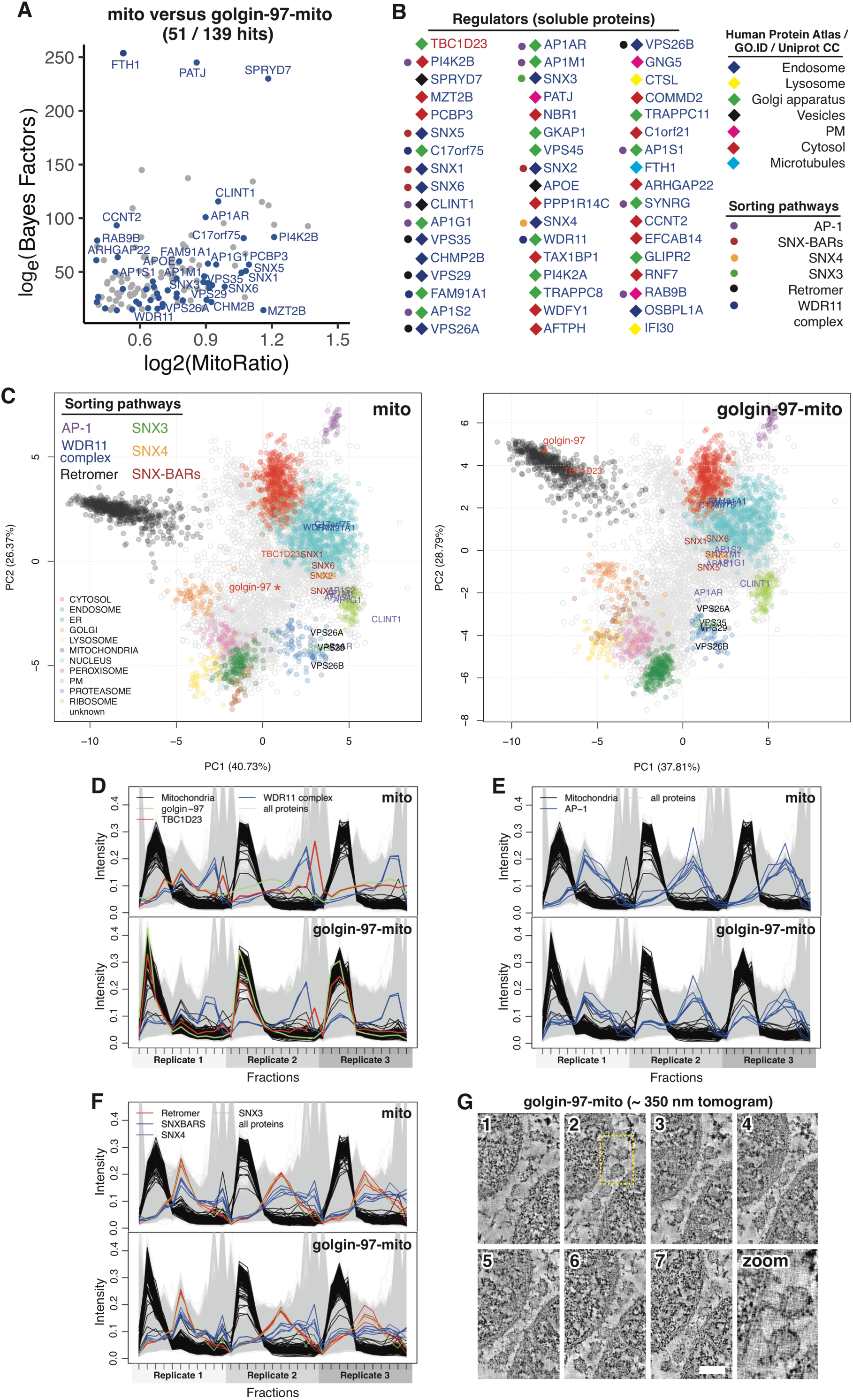
Adaptor proteins and accessory factors of vesicles specific to golgin-97 and golgin-245. **A**. MitoRatio versus Bayes Factor analysis of mito versus golgin-97-mito showing all hits (blue) that are both localised to the endosomal network and are also soluble proteins. **B**. All hits shown in (A) ordered from left column to right column based on descending MitoRatio scores. Classification of their localisation and sorting pathway is based on multiple localisation databases and previous publications. **C**. PCA projections for the LOPIT-DC of mito versus golgin-97-mito showing SVM classifications and the indicated proteins coloured based on the sorting pathway they belong to (A,B). **D-F**. TMT reporter ion distributions of proteins across fractions for each replicate for mito versus golgin-97-mito showing the profiles of known mitochondrial markers (Mitochondria) and all hits from (A,B) that belong to the indicated sorting pathways. **G**. Electron tomogram of resin-embedded HeLa cells expressing Golgin-97-mito and TMEM87A-RFP throughout a thickness of approximately 350 nm (208 sections). Mitochondria are large dense structures sandwiching balls. Scale bars, 100 *µ*m.

An alternative explanation for our enrichment of adaptor proteins could be that golgin-97-mito may be recruiting endosomes or TGN ribbons to the mitochondria (6). Lysosomal hydrolases are key markers of anterograde transport that traffic through the TGN and endosomes as they work their way to the lysosome. We did not observe any noticeable shift of lysosomal hydrolases to the mitochondria by golgin-97-mito in our analysis (Fig. S3A). We also performed electron tomography on golgin-97-mito expressing cells and found a striking accumulation of spherical structures surrounding the mitochondria that were roughly 30 to 100 nm in diameter (Fig. 3G & Movie S1). Together, this data firmly supports a model that golgin-97-mito captures AP-1, SNX3, SNX4 and SNX-BAR-derived retrograde transport vesicles. It also suggests that a subset of adaptor proteins remain on endosome-to-Golgi vesicles as they voyage to the TGN.

### Redirection of peroxisomes by mitochondrial golgins

The ectopic tethering of large numbers of vesicles to golgin-coated mitochondria is unlikely to have no indirect effects on the rest of the cell. This is evident by the drastic change in morphology of the mitochondria by electron microscopy as they become increasingly zippered together by vesicles (1). We detected a total of 43 proteins affected by golgin-97-mito that did not belong to the endosomal network (Fig. 4A,B). Of these, 34 were peroxisome proteins which were shifted to a more mitochondrial profile by both golgin-97-mito and GCC88-mito (Fig. 4C). The extent of this shift was more pronounced with the former, which correlated with its stronger overall ability to capture vesicles (Fig. 1G). Consistent with this, peroxisomes were significantly more adjacent to mitochondria in the presence of golgin-97-mito when localised by immunofluorescence (Fig. 4D). However, mutation of the *trans*-Golgi golgins did not affect peroxisome localisation (data not shown), indicating that the golgins do not directly affect peroxisomes themselves. Peroxisomes and mitochondria have a co-dependent relationship in the β-oxidation of fatty acids and the detoxification of reactive oxygen species (30). Moreover, evidence of peroxisome-mitochondria contact sites, particularly at mitochondria-ER junctions, have been reported in yeast, along with peroxisomal membrane protrusions that interact with mitochondria in mammalian cells (31-33). Our LOPIT-DC data provides further evidence of this co-dependence in response to what may be an unusual form of mitochondrial stress in mammalian cells.

**Figure 4.**
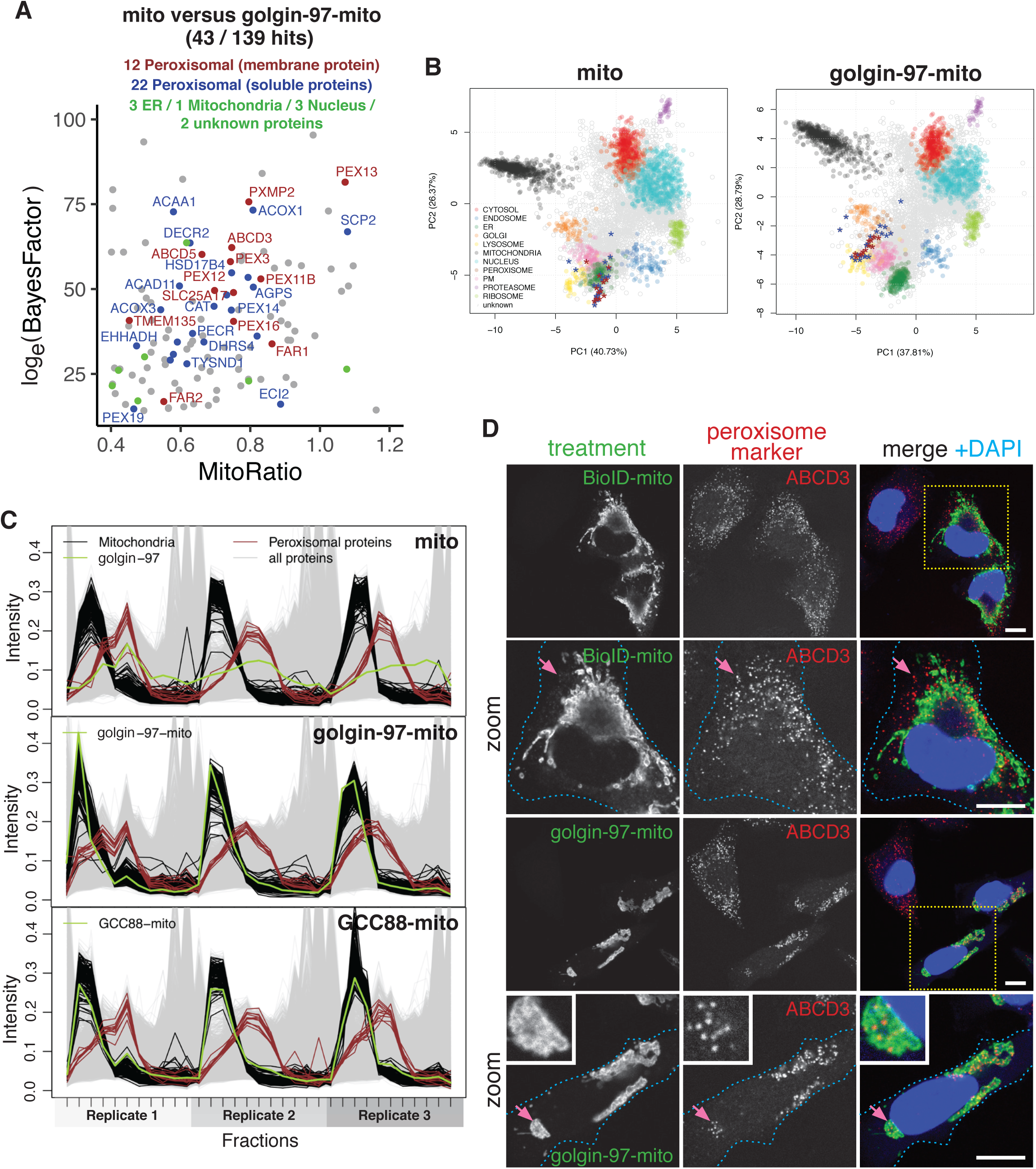
Redirection of peroxisomes by mitochondrial golgins. **A**. MitoRatio versus Bayes Factor analysis of mito versus golgin-97-mito showing all hits (brown, blue and green) that are not localised to the endosomal network. Classification of their localisation is indicated and is based on multiple localisation databases. **B**. PCA projections for the LOPIT-DC of mito versus golgin-97-mito showing SVM classifications along with peroxisomal membrane proteins (brown) and peroxisomal soluble proteins (blue) from (A). **C**. TMT reporter ion distributions of proteins across fractions for each replicate for mito, golgin-97-mito and GCC88-mito showing the profiles of known mitochondrial markers (Mitochondria) and peroxisomal membrane protein hits (A,B) (Peroxisomal Proteins). **D**. Confocal micrographs of HeLa cells expressing BioID-mito and the indicated golgin-mito construct (HA) stained for endogenous ABCD3 (peroxisome marker) and with DAPI. Scale bars, 10 *µ*m.

### GCC88 and golgin-97 share a distinct pool of TMEM87A and TGN46 enriched vesicles

TMEM87A is a member of the enigmatic LU7TM family of GPCR-related proteins that has been implicated in endosome-to-Golgi retrograde transport and that localises to the TGN (4, 34, 35). Of 16 total proteins, TMEM87A and TGN46 were distinctly affected by GCC88-mito (Fig. 1F). These proteins shared nearly identical profiles in the control and both were strikingly shifted toward the mitochondria by GCC88-mito (Fig. 5A-C), indicating that their localisation to membranes may be closely linked. As with TGN46, TMEM87A was also relocated to the mitochondria by golgin-97-mito (Fig. 5B,C & Fig. 2B). Furthermore, TMEM87A-GFP localised to spherical structures of roughly 50 nm in size tethered adjacent to the mitochondria by golgin-97-mito with correlative light electron tomography (CLEM) (Fig. 5D & Fig. 3G) (36). Thus, TMEM87A is enriched in distinct retrograde transport vesicles.

**Figure 5.**
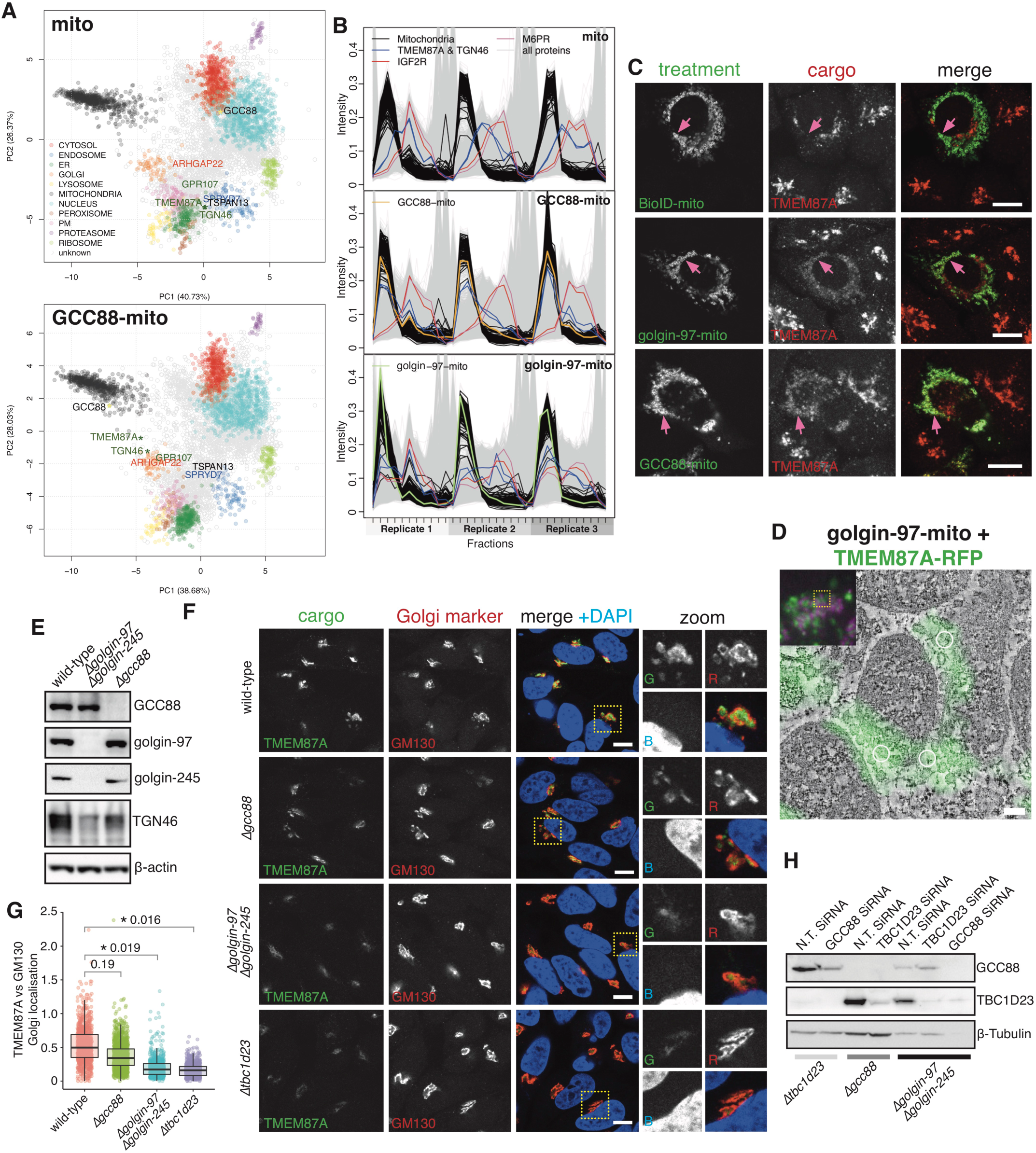
GCC88 and golgin-97 share a distinct pool of TMEM87A and TGN46 enriched vesicles. **A**. PCA projections for the LOPIT-DC of mito versus GCC88-mito showing SVM classifications and top 6 hits that are localised to the endosomal network from Fig. 1F. **B**. TMT reporter ion distributions of proteins across fractions for each replicate for mito, golgin-97-mito and GCC88-mito showing the profiles of known mitochondrial markers (Mitochondria) and the indicated proteins. **C**. Confocal micrographs of HeLa cells expressing BioID-mito the indicated golgin-mito construct (HA) stained for endogenous TMEM87A (cargo). Scale bar, 10 *µ*m. **D**. CLEM using super-resolution fluorescence microscopy (FM) of resin-embedded HeLa cells expressing Golgin-97-mito and TMEM87A-RFP (green). Open white circles indicate location of a fiducial marker for correlation. Inset shows the FM image (yellow box) emitting TMEM87A-RFP (Green) and mitotracker (purple) that was used for correlation; scale bar, 100 *µ*m. **E**. Immunoblot of proteins from wild-type and mutant HeLa cells. **F**. Confocal micrographs of wild-type and mutant HeLa cells stained for endogenous TMEM87A (cargo), GM130 (marker of *cis*-Golgi) and with DAPI. Scale bar, 10 *µ*m. **G**. Quantification of (F) measuring the ratio of the mean intensity of TMEM87A at the Golgi over the intensity of GM130 at the Golgi (Golgi localisation). Boxplots are of Golgi localisation of Golgi from a minimum of 300 cells across 3 independent replicates. p-values are of two-sample t-tests comparing the overall mean Golgi localisation of each replicate. **H**. Immunoblot of proteins from wild-type and mutant HeLa cells transfected with the indicated SiRNA over the course of at least a week.

The retrograde trafficking of IGF2R and M6PR from endosomes to the TGN involve clathrin/AP-1, retromer-dependent and independent pathways (15, 26, 27, 37). Strikingly, our LOPIT-DC of GCC88-mito had no significant effect on either of these cargo (Fig. 5B), suggesting that the distinct pool of TMEM87A and TGN46 enriched vesicles that GCC88 captures is not dependent on these particular retrograde pathways. GCC88-mito has previously been reported to capture IGF2R and M6PR enriched vesicles when expressed by transient transfection (1, 7, 37). It is likely that our stable expression of GCC88-mito used in our LOPIT-DC experiments were below this threshold, allowing us to only detect one distinct pool of vesicles.

TGN46 has previously been shown to depend on golgin-97, golgin-245 and TBC1D23 for Golgi recruitment (8). To extend this study to GCC88, we generated *Δgcc88* mutants using CRISPR-Cas9 (Fig. 5E). When TGN46 is missorted from endosomes, it is diverted to lysosomes and degraded, allowing efficacy of its sorting to be quantified by both immunofluorescence and blotting (38). We did not observe any noticeable defect in TGN46 localisation in the *Δgcc88* mutant by immunofluorescence (data not shown), and observed only a mild decrease in TGN46 levels by blotting (Fig. 5E). Similarly, the localisation of TMEM87A to the TGN became significantly more diffuse in *Δtbc1d23* mutants and in *Δgolgin-97 / Δgolgin-245* mutants by immunofluorescence, whereas *Δgcc88* mutants did not significantly affect Golgi localisation (Fig. 5F,G). Taken together, these results suggest that golgin-97 and golgin-245 also recruit TMEM87A and TGN46 enriched vesicles via TBC1D23, and are much more efficient at capturing these vesicles than GCC88.

Our results clearly indicate that GCC88 is redundant with golgin-97 and golgin-245 in capturing certain pools of vesicles. In firm support of this, knockdown of GCC88 in *Δgolgin-97 / Δgolgin-245* mutants prevented cells from growing (Fig. 5G). Importantly, the combined loss of TBC1D23 and GCC88 did not affect cell growth. These results clearly show there are TBC1D23-independent mechanisms that allow golgin-97 and golgin-245 to capture vesicles which likely play a more prominent role in Golgi tethering. Our LOPIT-DC of golgin-97-mito using our Bayesian analysis has illustrated the power of this approach to organelle proteomics, and its application to cells expressing mitochondrially localised golgins has demonstrated that this is an effective means of characterising the cargo of intracellular transport vesicles. The study thus provides an extensive list of cargo proteins that move in these retrograde routes, as well as peripheral membrane proteins that may act on these carriers and hence will hopefully be a useful resource for future studies on this important aspect of intracellular membrane traffic.

## Author Contributions

J.J.H.S and S.M. devised and J.J.H.S, S.M. and K.S.L. planned the study. J.J.H.S performed the LOPIT-DC experiments. J.J.H.S and S.Y.P.C performed the mass spectrometry. J.J.H.S and O.M.C. analysed the LOPIT-DC data. J.J.H.S. and J.C.O performed the immunofluorescence and blotting experiments. A.B. and J.C. performed the electron tomography. J.J.H.S and S.M. wrote the manuscript.

## Acknowledgments

We are indebted to A. Gillingham for comments on the manuscript, to J.A. Christopher for her technical assistance and aid with LOPIT-DC, to O.L. Vennard for help in multimedia used in figures, to Mike Deery and Yagnesh Umrania of the Cambridge Centre for Proteomics, and to all the members of the Medical Research Council (MRC) Laboratory of Molecular Biology Mass Spectrometry Unit. Funding was made from the MRC (MRC file reference number MC_U105178783), and by a European Molecular Biology Organisation long-term fellowship to J.J.H.S.

## Figure Legends

**Supplementary Figure 1.**
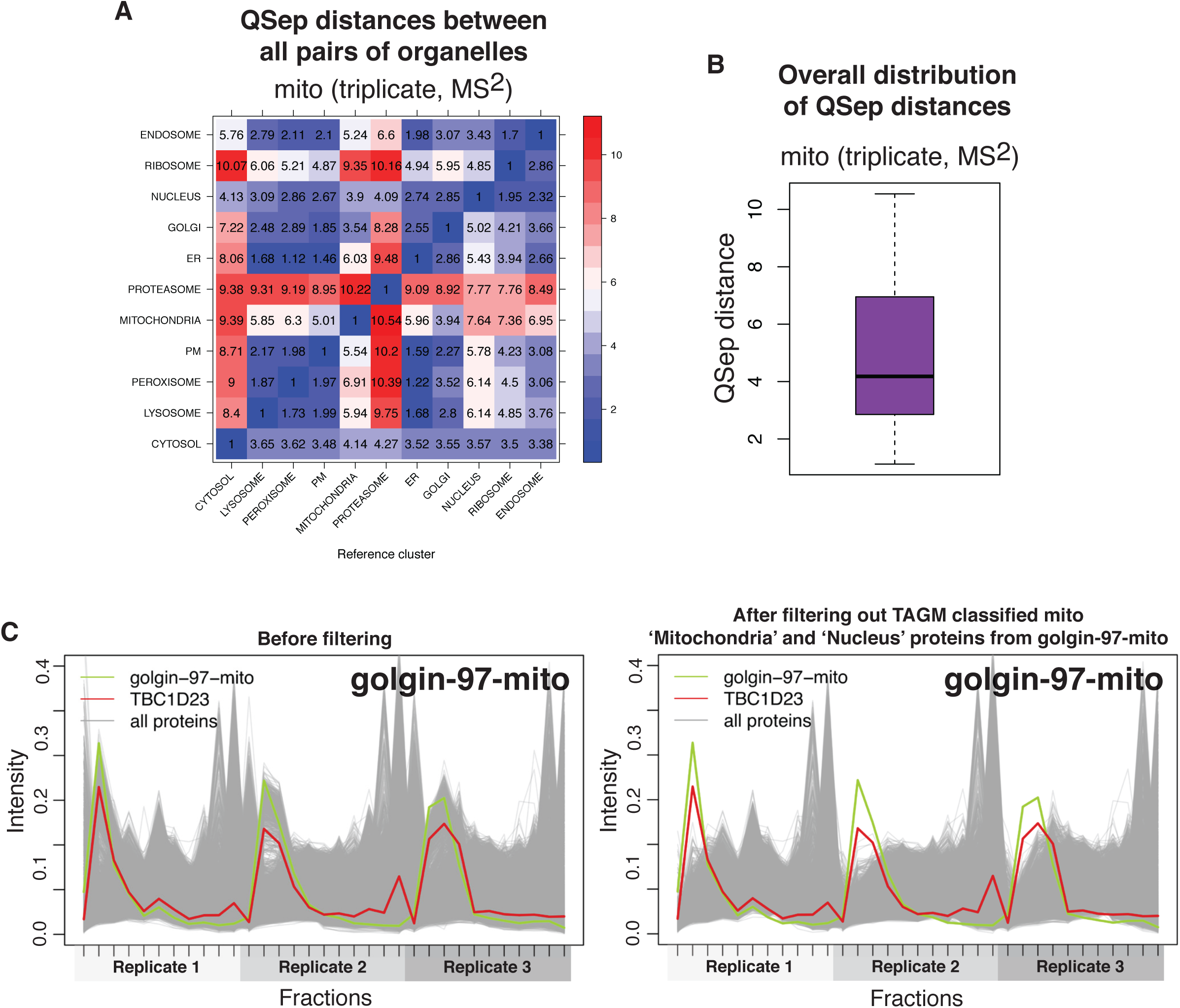
Assessing and pre-filtering LOPIT-DC data. **A**. QSep distance quantifying separation between all pairs of compartments in the LOPIT-DC of mito. **B**. The overall distributions of QSep distances in the LOPIT-DC of mito. These QSep results should be compared to (2). **C**. TMT reporter ion distributions of proteins across fractions for each replicate for golgin-97-mito showing the effect of discarding TAGM classified ‘Mitochondria’ and ‘Nucleus’ proteins from mito in golgin-97-mito.

**Supplementary Figure 2.**
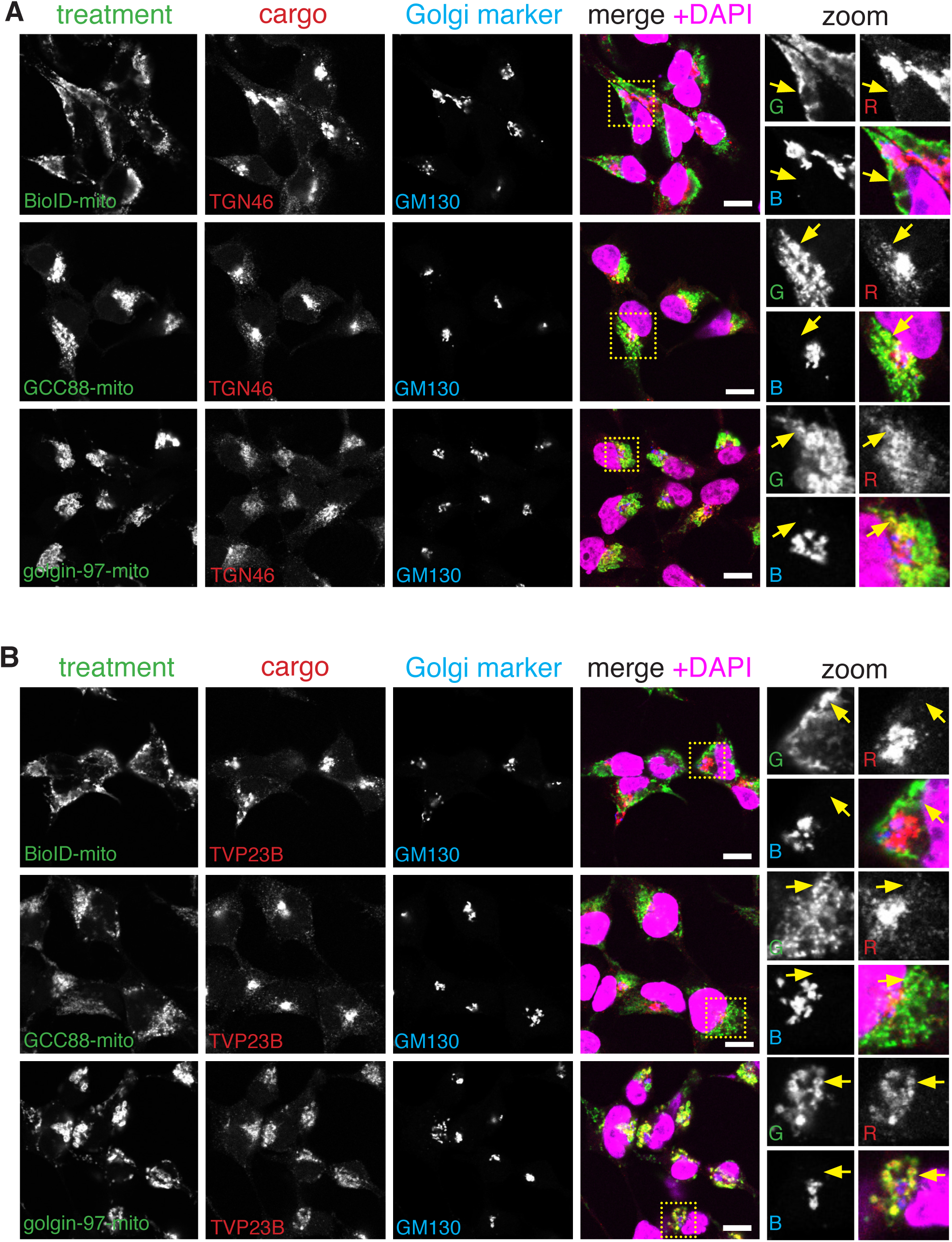
Mito-relocation of cargo by golgin-97-mito and GCC88-mito. **A,B**. Confocal micrographs of 293T cells stably expressing the doxycycline-inducible BioID-mito, golgin-97-mito and GCC88-mito constructs (HA) stained for endogenous TGN46 or TVP23B (cargo), and GM130 (marker of *cis*-Golgi) and with DAPI. Cells were incubated with 1 *µ*g ml^−1^ doxycycline for 48 h. Scale bar, 10 *µ*m.

**Supplementary Figure 3.**
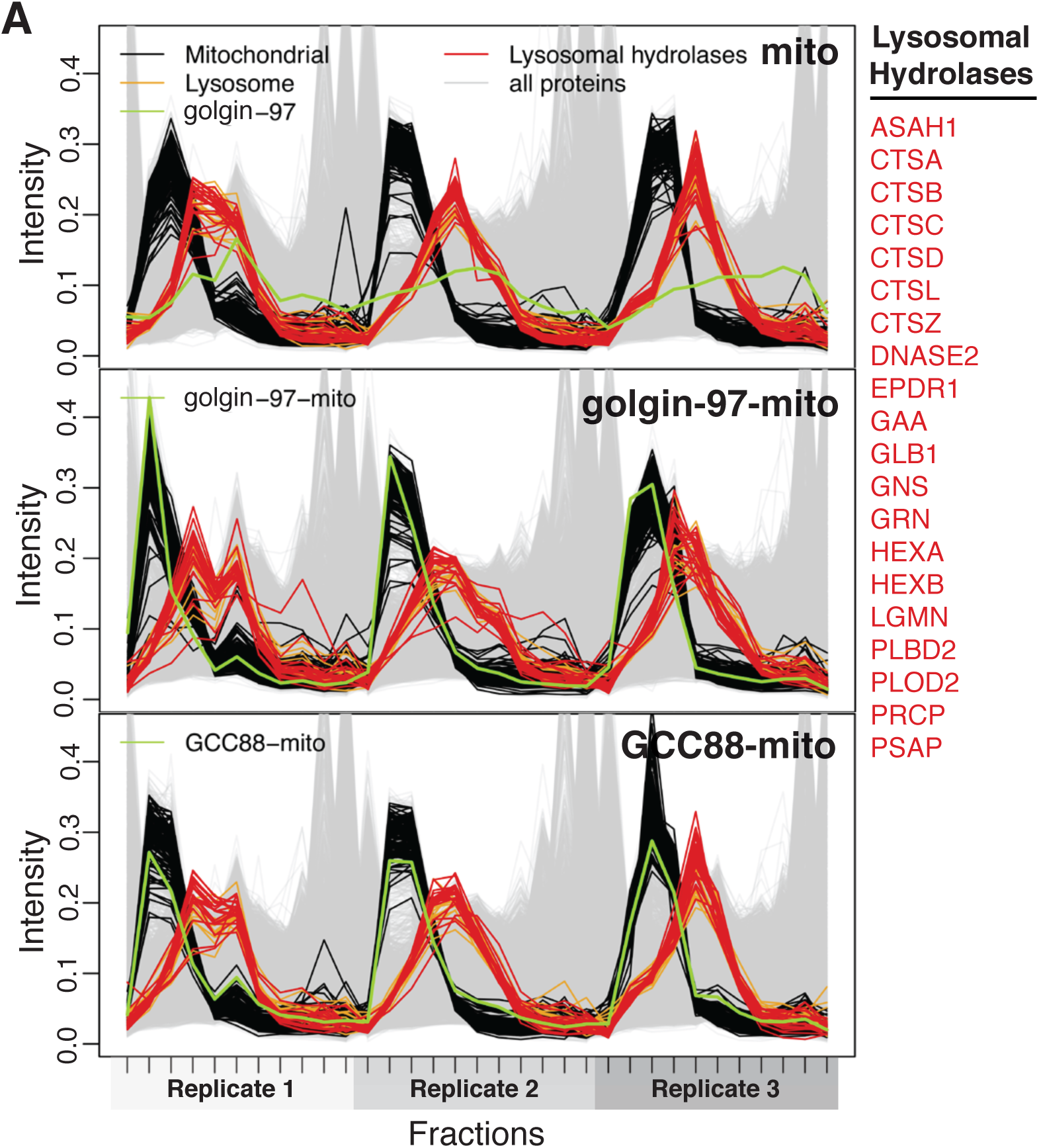
Lysosomal hydrolases are not shifted to a mitochondrial profile by golgin-97-mito or GCC88-mito. **A**. TMT reporter ion distributions of proteins across fractions for each replicate for mito, golgin-97-mito and GCC88-mito showing the profiles of known mitochondrial markers (Mitochondria), known lysosomal markers (Lysosome) and lysosomal hydrolases that have previously been shown to be found on anterograde Golgi-to-endosome clathrin/AP-1 vesicles (15-17).

## Methods

### Plasmids and Antibodies

The golgin-mito plasmids used for transient transfection have been previously described (1, 7, 8). These constructs were subcloned into pcDNA5/FRT/TO to make stable lines expressing golgin-97-mito (pMX0267), GCC88-mito (pmX0269), along with mito (HA tagged to the monoamine oxidase (MAO) transmembrane domain; pMX0266). Furin-GFP, Furin (Human CCDS10364.1) tagged to EGFP in pcDNA3.1+ (pJJS084); TMEM87A-RFP, TMEM87A cloned from 293 cells tagged to mRFP in pcDNA3.1+ (pJCO003); BioID-mito, BioID with an HA tag linked to the MAO transmembrane domain in pcDNA3.1+ (pJJS112) and pcDNA5/FRT/TO (pJJS128). CRISPR-Cas9 used the plasmids eSpCas9 (Addgene no. 71814) and pIRESpuro3 (Clontech). eSpCas9 was digested with BbsI and the following gRNA sequences were inserted by annealed oligonucleotide cloning: 5’-TCAGCAGGCTTTTATAGGCA -3’ targeting human GCC88 (pJJS294). All antibodies used in this study have previously been described in (8) with the exception of the following: ATG9A, Abcam - ab108338; TVP23B, Human Protein Atlas - HPA019585; TMEM87A, Human Protein Atlas - HPA018104; ABCD3, Human Protein Atlas - AMAb90995; TBC1D23, Proteintech 17002-1-AP; Alpha-tubulin, Rat monoclonal to alpha-tubulin clone YL1-2 made in house.

### Cell culture, transfection, CRISPR knockout cell lines and immunofluorescence

Cell lines HeLa (ATCC) and 293T Flp-In T-REx (ThermoFisher Scientific, R78007) were cultured in Dulbecco’s modified Eagle’s medium (DMEM; Invitrogen) supplemented with 10% fetal calf serum (FCS) and penicillin / streptomycin at 37 °C and 5% CO_2_. HEK 293T Flp-In T-REx cell lines were transduced with pMX0267, pmX0269, pMX0266 and pJJS128 as previously described (8). The *Δtbc1d23* mutant and *Δgolgin-97 / Δgolgin-245* mutant was previously described in (8) and *Δgcc88* was generated using the same protocol with pJJS294. All cell lines were tested regularly to ensure that they were mycoplasma free (MycoAlert, Lonza). For immunofluorescence, cells were transfected with plasmid DNA using FuGENE 6 according to the manufacturer’s instructions (Promega). Cells were fixed with 4% formaldehyde in PBS and permeabilised in 0.5% (v/v) Triton X-100 in PBS. Cells were blocked for one hour in PBS containing 20% FCS and 0.25% Tween-20. If this protocol did now show good Golgi staining for a particular antibody, cells were permeabilised and blocked in 20% FCS and 0.1% saponin. Cells were probed in the same buffer as the primary and secondary antibodies. Each antibody used in immunofluorescence was diluted to a maximum concentration of 100X. All immunofluorescence experiments were performed at a minimum of three times. For quantification of MitoRatio, cells and the mitochondria and Golgi were defined with DAPI, HA to the mito construct and GM130, respectively (NIS-Elements, Nikon). Then the mean intensity of the cargo channel in the mitochondria mask (excluding the areas they combine with the Golgi mask) were divided by the mean intensity of the cargo channel of the Golgi mask for each quantified cell. The mitochondria and Golgi mask had a minimum threshold of 18% if the mean labelling of their respective markers to exclude poorly expressed constructs or out-of-focus Golgi. For quantification of Golgi localisation, the mean intensity of the cargo in the Golgi mask was divided by the mean intensity of GM130 in the Golgi mask. The same minimum threshold was used to exclude out-of-focus Golgi.

### siRNA knockdown of HeLa cells

On Day 0, equal numbers of *Δtbc1d23, Δgcc88* and *Δgolgin-97 / Δgolgin-245* HeLa cells were seeded in a 24-well plate. Cells wells were transfected on day 1 and 4 using RNAiMax (Invitrogen) with either non-targeting SiRNA (ON-TARGETplus control D-001810-01-05), SiRNA to TBC1D23 (Dharmacon ON-TARGETplus human TBC1D23 SMARTpool 55773), or SiRNA to GCC88 (Dharmacon ON-TARGETplus human GCC1 SMARTpool L-017478-00-005) according to manufacturer’s instructions. On day 8, cells were washed 2x with PBS and collected with 1x LDS sample buffer and solubilized by sonication. Samples were analysed by western blotting.

### Immunoblotting

One 75 cm^2^ flask of cells at ∼90% confluence was harvested by scraping and centrifugation, washed twice with ice-cold PBS, resuspended in 30 to 500 *µ*l lysis buffer (50 mM Tris pH 7.4, 0.1 M NACl, 1 mM EDTA, 1% Triton X-100, 1 mM PMSF, cOmplete inhibitors) for 30 min on ice, clarified by centrifugation, and protein concentration was determined (BCA protein assay - ThermoFischer). Lysate concentrations were normalised, and the lysates were boiled in NuPAGE SDS sample buffer containing 10% β-mercaptoethanol at 90 °C for 5 min, run on a gel and transferred to nitrocellulose. All blots were blocked in 5% (w/v) milk in PBS-T (PBS with 0.1% (v/v) Tween-20) for 1 h, incubated overnight at 4 °C with primary antibody in the same blocking solution, washed three times with PBS-T for 5 min, incubated with HRP-conjugated secondary antibody in 0.1% (w/v) milk in PBS-T for 1 h, washed five times with PBS, and detected with Immobilon Western HRP substrate. All antibodies were diluted to a concentration of a minimum of 1000X. All immunoblotting experiments were performed at a minimum of two times.

### Correlative FM and electron tomography of resin-embedded cells

HeLa Cells grown on 3 mm sapphire disks (Engineering Office M. Wohlwend, Switzerland) in six-well plates for 24 hr, transfected with 1000 ng golgin-97-mito and 1000 ng TMEM87A-RFP using FuGENE 6 for 24 hr, and then high-pressure frozen using a HPM100 (Leica Microsystems). The remaining protocol was performed as (36).

#### LOPIT-DC

LOPIT-DC was performed as (2) with the following exceptions. 293T cells were passaged in at least 8X T175 flasks per treatment in medium so that they were approximately 50% confluent after 24 hr. After this time, 1 *µ*g ml^−1^ doxycycline was added to the medium and incubated for 48 h. The cells were then collected through gentle pipetting and lysed as in (2). The remaining cell debris from 3X 20 min 200 g spins of the cell homogenate were collected and used in a nuclear prep (OptiPrep Application Sheet S10) that became an additional TMT-labelled fraction (Fig. 1B). The concatenated RP-UPLC pre-fractionated TMT-labelled peptides were run on a Q-Exactive Plus Orbitrap LC-MS/MS system and subsequent analysis of LOPIT-DC data was optimised for MS^2^ as in (11). All LOPIT-DC experiments were performed at a minimum of three independent times.

#### Posterior localisation probabilities

The posterior probability that a protein belongs to a sub-cellular niche, henceforth referred to as the posterior localisation probability, is computed using the TAGM-MAP method (3). Briefly, the parameters of the T-augmented Gaussian mixture model are learnt by maximising the log posterior of the parameters with respect to the data, to obtain *maximum a posteriori* (MAP) estimates of the parameters. The posterior localisation probability that a protein belongs to each organelle is then computed and the most probable sub-cellular niche is reported. Default prior choices are made as described in (3) and (39). Proteins that are predicted to localise to the mitochondria and nucleus are discarded from subsequent analysis.

#### Bayesian non-parametric two sample test

To detect perturbations in the quantitative protein profiles, we apply a Bayesian non-parametric two sample test (40). First, the data are transformed using the additive log ratio transform (41). We then proceed to test whether the protein profiles are different between the control and treatment. Formally, we test against two contrasting models. The first model posits that the quantitative protein profiles in each experiment (control and treatment) are drawn from an identical shared distribution. Whilst the second supposes that there are independent models for each of the control and treatment. The log Bayes factor is used to objectively determine support for one model over the other, where larger log Bayes factors are considered support for the independent model (42). A non-parametric prior over functions, the Gaussian process, is specified with squared exponential covariance (43). The hyperparameters for the Gaussian process are given the default Gamma priors as specified in (40). The natural logarithm of the Bayes factors is reported.

#### Mitochondrial Ratios

To determine the biological relevance of quantitative protein profiles that are shifted between treatment and control, we compute a mitochondrial ratio (MitoRatio) as follows. For each protein in each experiment, the squared Mahalanobis distance (44) to the mean of the quantitative profiles of the mitochondrial marker proteins is computed, where a robust estimate for the covariance is used (45). Then, for each protein, the ratio of the distances in control and treatment is computed, proceeded by a log2 transform. Proteins that move closer to the mitochondria upon the treatment have larger mitochondrial ratios.

